# Single-cell Organelle Extraction with Cellular Indexing

**DOI:** 10.1101/2024.12.23.630180

**Authors:** Trinh Lam, Alison Su, Ana E. Gomez Martinez, Anna Fomitcheva-Khartchenko, Amy E. Herr

## Abstract

Bulk methods to fractionate organelles lack the resolution to capture single-cell heterogeneity. While microfluidic approaches attempt to fractionate organelles at the cellular level, they fail to map each organelle back to its cell of origin—crucial for multiomics applications. To address this, we developed VacTrap, a high-throughput microfluidic device for isolating and spatially indexing single nuclei from mammalian cells. VacTrap consists of three aligned layers: (1) a Bis-gel microwells layer with a ‘trapdoors’ (BAC-gel) base, fabricated atop a through-hole glass slide; (2) a PDMS microwell layer to receive transferred nuclei; and (3) a vacuum manifold. VacTrap operation begins with cell lysis using DDF to release intact nuclei into the Bis-gel microwells, while cytoplasmic proteins are electrophoresed into the Bis-gel. Upon exposure to DTT and vacuum force, the trapdoors open, allowing nuclei to transfer to the PDMS microwells. VacTrap dissolves the trapdoors within 3-5 minutes and achieve synchronized nuclei transfer with 98% efficiency across 80% of trapdoors in a 256-microwell array, surpassing the <1% efficiency of passive transfer without vacuum. Morphology analysis confirmed preservation of organelle integrity throughout VacTrap operation. By enabling spatial indexing of nuclei back to their original cell, VacTrap provides a robust, high-throughput tool for single-cell multiomics applications.

## Introduction

Cells are composed of specialized organelles that each perform unique functions. Structural and functional assays rely on organelle isolation, wherein the integrity of the isolated organelles directly affects analysis accuracy, ultimately shaping our understanding of biology [1]. Bulk organelle-fractionation methods (e.g., density-gradient centrifugation, immune-isolation, free-flow electrophoresis, detergent-based chemical fractionation, enzymatic digestion) are labor-intensive, designed for pooled cell suspensions and not suitable for sparingly available specimens, and offer low organelle- recovery yields. Although suffering from these performance shortcomings, density-gradient centrifugation remains widely used [2–4]. Immuno-isolation is constrained by the availability and quality of antibody probes specific to organelle surface proteins [5]. Free-flow electrophoresis separates cellular organelles [6, 7] with low recovery purity and resolution [8]. Detergent cocktails enrich specific cellular fractions; with each chemical component having a distinct solubilization efficiency [9, 10]. Yet, enzymatic treatments are known to perturb cell-cycle status, apoptosis, and structural alterations [11, 12]. Overall, bulk organelle-isolation methods require multiple steps requiring extensive manual handling and yield compromised purity and integrity of the isolated organelles, thus impacting functional analysis.

Microfluidic technologies offer enhanced precision in organelle isolation from sparingly available starting samples and can overcome limitations of isolated-organelle yield and sample-prep throughput. These tools include techniques utilizing magnetic nanoparticles [13], immuno-affinity [14, 15], flow-based or channel structures [16–19], digital microfluidics [20, 21], magnetophoretic-based microfluidics [22], and devices structured to capture DNA [23, 24]. While precise, multi-step process flows (e.g., on-chip extraction, isolation, and off-chip recovery) can be a source of organelle damage and yield loss.

Even with the advent of precision tools for organelle isolation, the post-isolation pooling of isolated organelles remains ubiquitous, making even contemporary microfluidic techniques incompatible with follow-on single-cell or single-organelle analyses that require indexing of organelle back to the originating cell. Indexing an isolated organelle to the originating cell forms a basis for understanding organelle-derived heterogeneity that exists between cell types and among individual cells, even of the same type [25]. In a related aspect of performance: the preservation of spatial information is increasingly sought, such as mapping an isolated organelle(s) back to the originating tissue context. Logically, mapping back to the single originating cell is also sought because functional links between biological processes can (and do) occur at the level of single cells.

An active area of organelle- and cellular-level biology is the study of the nucleus as a coordinating – and typically the largest – cellular organelle. Microfluidic tools make single-nucleus measurements possible. To analyze chromosomal DNA from single nuclei, Benítez et al. introduced a micropillar array and hydrodynamic flows to extract and stretch chromosomal DNA from ∼100 single mammalian cells per chip [26]. Following these precise, in-situ assays of chromosomal-DNA stretching from a single cell, DNA was recovered and quantified off chip. Similarly, Wang et al. utilized microchannel geometries to isolate and stretch chromosomal DNA from 10-20 nuclei for subsequent DNA fluorescence in-situ hybridization (FISH), with each signal traced back to its originating nucleus [27]. While limited to assessing DNA damage, conventional agarose-slab embedded and microwell-based comet assays, do allow researchers to assess DNA damage and map back to originating cell [28]. These existing techniques point to the promise for assessing other nuclear components -- including proteins such as transcription factors – and mapping said measurements back to the originating cell and/or tissue context.

With a focus on introducing tools for single-cell resolution protein measurement, our group introduced a suite of single-cell immunoblotting modalities designed using microwell- isolated single cells, including single-cell western blots [29–31]. With an eye towards organelle-biology and subcellular omics, we introduce tools for single-nucleus isolation using microwell-isolated mammalian cells subjected to differential detergent fractionation (DDF), a technique that employs a sequence of detergents with varying solubilization strengths to selectively extract and separate cellular components based on membrane properties [2, 32]. In one example of a single-nucleus resolution analysis made possible by combining microfluidic precision with DDF, we performed a single-cell Western blot of each cytoplasmic compartment and a distinct electrophoresis of each nuclear compartment for an array of cells. In a second example that extends on the assay just described, the Western blot analysis of each single nucleus was swapped out with a PCR assay, allowing both cytoplasmic protein targets and nuclear DNA and RNA targets to be detected in the same originating cell [33–35]. While both single-nucleus precision assays are suitable for sparingly available starting specimens (<10 starting cells, e.g., isolated circulating tumor cells, individual blastomeres comprising two- and four-cell preimplantation murine embryos), sample and analysis throughput must be increased for applicability to larger-cell-number specimens.

Here, we introduce a single-cell resolution organelle isolation method incorporating a single-cell isolation via a polyacrylamide microwell array that is optimized for nuclear isolation after DDF. We utilize microfluidic automation to enhance throughput while offering the capacity to isolate and then index individual nuclei back to each originating cell. In a multi-layered, planar microfluidic device, individual cells are isolated by settling into an array of polyacrylamide microwells, one cell per microwell. Perturbation, proteomics, or imaging analysis can be performed on intact cells in these top-layer polyacrylamide gel microwells. To isolate and extract single nuclei for further analysis, each cell’s cytoplasmic membrane is lysed using DDF, and one intact nucleus remains in each microwell. To concurrently transfer each nucleus to an aligned PDMS microwell situated below the PAG microwell, an interleaving layer of through holes filled with a dissolvable gel is actuated. These dissolvable ‘trapdoor’ in the floor of each PAG microwell opens when the dissolvable gel is exposed to reducing agents (i.e., dithiothreitol (DTT)) and suction is applied using an attached microfluidic vacuum manifold. Once the trapdoors are open, 100’s of nuclei are simultaneously transferred from the PAG microwell array to the PDMS microwell array, at one nucleus per microwell occupancy. Here, we detail the multi-layer fluidic design, chemical and hydrodynamic control optimization, and resultant organelle isolation and extraction performance of this single-nucleus isolation and extraction technique.

## Results and Discussion

To advance organelle biology and sub-cellular omics, our study introduces a device to extract nuclei from 100’s of individual mammalian cells using microfluidic automation, precision handling, and subsequent indexing of intact nuclei back to the originating cell (**Figure 1**). The multilayer microfluidic device (called VacTrap, for brevity) is designed to perform a controlled, automated single nucleus preparation protocol. We report here on device design and fabrication, optimization of the chemical and mechanical functions of VacTrap (device and preparation protocol), and performance of the single-nucleus extraction system.

**Figure 1.**
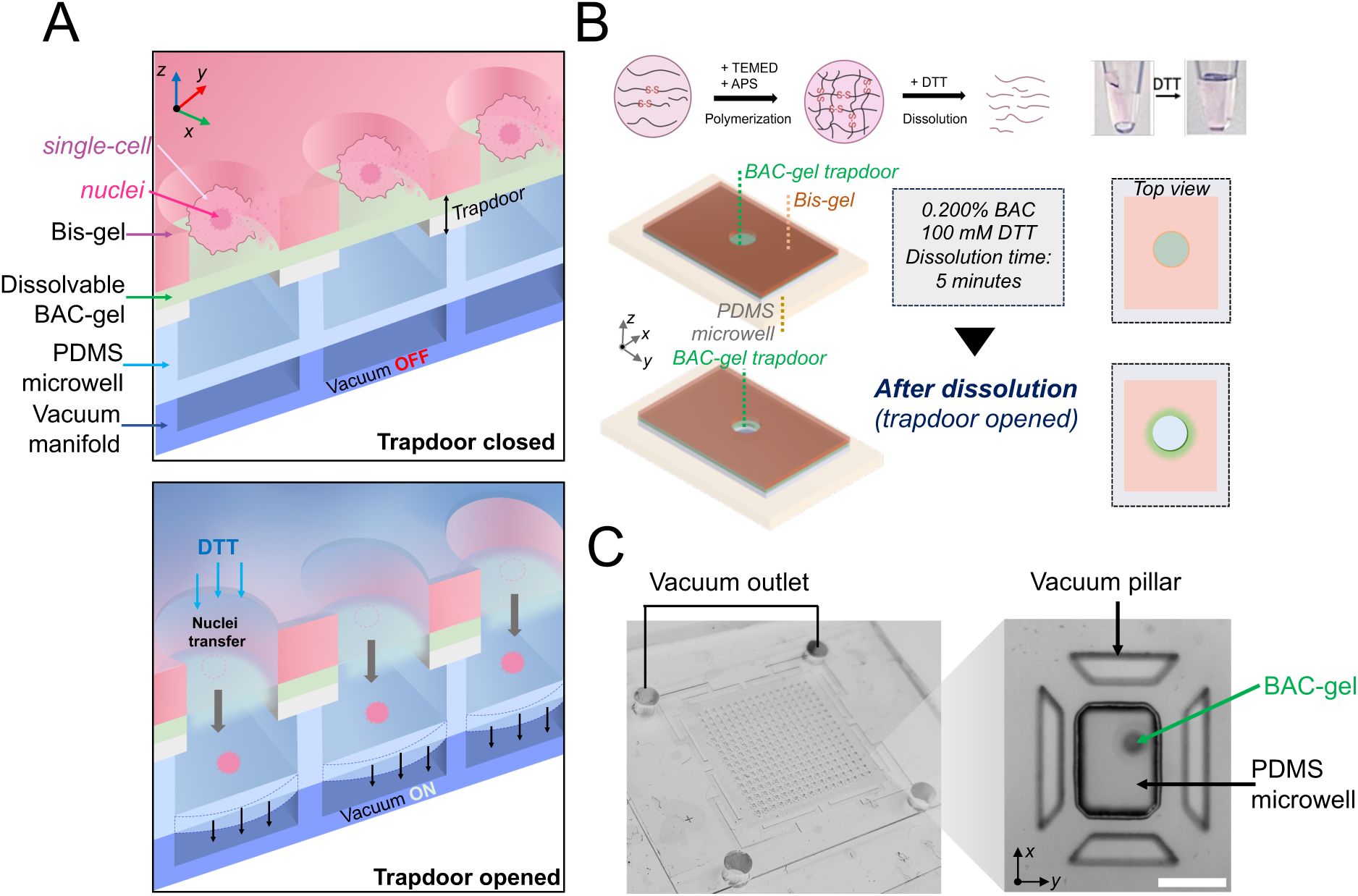
Single-nucleus extraction with cellular indexing uses a co-planar multilayer microfluidic device with vacuum-assisted actuation, called VacTrap. (A) Schematic of VacTrap which comprises three co-planar device layers: (1) a whole-cell receiving layer which is the top, open-fluidic Bis-gel microwell layer (purple arrow), with each of 256 microwells having a mechano-chemically actuated BAC-gel ‘trapdoor’ at the base (green arrow), (2) a nucleus-receiving layer molded with a PDMS microwell array (blue arrow), wherein each PDMS microwell is aligned to an upper Bis-gel microwell and BAC-gel trapdoor, and (3) a vacuum manifold layer to apply a suction force to drive the simultaneous transfer of nuclei across from the Bis-gel microwells to the receiving PDMS microwells. The mechano-chemically actuated trapdoor comprises a layer of reversibly crosslinked BAC hydrogel on a glass slide engraved with precision-drilled 100-μm diameter through holes, each aligned to a stacked microwell pair. To open the trapdoor, application of DTT depolymerizes the disulfide-cross-linked BAC-gel through a thiol–disulfide exchange reaction. Once liquified, application of suction from the vacuum manifold deflects the thin base membrane of the PDMS microwell layer, thereby creating a pulling action transfer a bolus of dissolved BAC-gel and isolated nucleus through each through hole and into the receiving PDMS microwell. (B) Schematic of the synthesis and dissolution mechanism of the dissolvable polyacrylamide BAC-gel used to create and open the trapdoor feature. Acrylamide monomers and BAC crosslinker undergo polymerization via C=C double bonds facilitated by ammonium persulfate (APS) and tetramethylethylenediamine (TEMED), resulting in the formation of a dissolvable polyacrylamide gel layer. Exposure to reducing agents (DTT) depolymerize the disulfide-crosslinked BAC-gel due to the thiol–disulfide exchange reaction. Upon dissolution with DTT and application of a suction force, the through hole is opened and materials transfer from the top Bis-gel microwell into the bottom receiving PDMS microwell. (C) Brightfield photograph of the assembled three-layer VacTrap assembly showing vacuum ports. Micrograph inset shows a top- down view of Bis-gel to PDMS microwell pair stack, interleaving trapdoor layer, and vacuum trapezoid pillars to prevent deformation of PDMS microwell during suction force applied.

### Overall design of the VacTrap nucleus isolation and extraction device

The VacTrap device design consists of three co-planar layers (**Figure 1A**): (1) a whole- cell receiving layer which is the top, open-fluidic, Bis-gel layer stippled with an array of microwells each having a mechano-chemically actuated ‘trapdoor’ at the base of each microwell. The trapdoor feature consists of a layer of chemically dissolvable BAC-gel coated on a glass support slide with through holes (**Figure 1B**), (2) a nucleus-receiving layer molded with a PDMS microwell array, wherein each PDMS microwell is aligned to an upper Bis-gel microwell and BAC-gel trapdoor, and (3) a vacuum manifold layer to apply a suction force that drives the simultaneous transfer of nuclei from the cell-laden Bis-gel microwells to the nucleus-receiving PDMS microwells (**Figure 1C**).

After sedimentation and imaging of intact whole cells in the Bis-gel microwells in the top whole-cell receiving layer, nuclei are concurrently isolated from the cells prior to transfer to the PDMS nucleus-receiving microwells. To isolate nuclei, we introduce a DDF buffer that selectively lyses each cell’s cytoplasmic membrane, leaving the nuclear membrane, and thus nucleus, intact in each top-layer Bis-gel microwell [31].

We selected three distinct materials for the whole-cell and nucleus-receiving microwell array layers: polyacrylamide gels (BAC-gel and Bis-gel), glass, and elastomer (PDMS).

First, the BAC-gel is polymerized atop a through-hole glass slide, creating a stable, covalently bonded trapdoor. After the BAC-gel polymerizes, the Bis-gel is then polymerized directly on top of the BAC-gel, forming the microwells with the BAC-gel as a base of the microwell. This layered assembly allows for efficient isolation of intact mammalian cells for downstream applications, such as perturbation, imaging, or proteomics analysis. The through-hole glass slide acts as a gate for isolating the nucleus from each cell, providing structural support throughout the single-cell handling steps and ensuring stability during the alignment to the PDMS nucleus-receiving microwells. Without the support of the glass, the thin composite of BAC and Bis-gels (∼100 um thick) would collapse during the dissolution process. Additionally, the through-hole glass slide forms a well-defined path for nuclei to travel from the Bis-gel microwells, through the trapdoor, and into the PDMS microwells. This setup physically transfers each nucleus into a PDMS compartment compatible of with standard biochemical processes, such as PCR. Glass is an ideal material for this design due to its strong bonding properties with both polyacrylamide and PDMS, which is commonly used in single-cell and molecule analyses. Its ability to form stable bonds with both polyacrylamide and PDMS makes it optimal for this system.

The trapdoor at the base of each top-layer Bis-gel microwell is designed to be initially closed, to open with chemical and mechanical triggers, and then remain open (**Figure 1A-B**, and **Figure 2A-B**). To achieve these functions, the trapdoor is composed of a layer of N,N’-bis(acryloyl)cystamine (BAC), a reversible crosslinker, polymerized with acrylamide monomer to form a dissolvable polyacrylamide gel layer (BAC-gel), cast on a 400-μm thick glass slide with laser-etched 100-μm diameter through holes (**Figure 2A and C**) [36]. Application of DTT results in the degradation of the disulfide-cross-linked BAC-gel due to the thiol–disulfide exchange reaction (**Figure 1B**) [37–39]. Application of a suction force to the bottom of the PDMS microwell receiving layer transfers force up to the sandwiched trapdoors and initiates nuclei transfer from the Bis-gel microwells into said PDMS receiving microwells. To be effective at transmitting the suction force from the bottom of the PDMS microwells to the trapdoors, the receiving PDMS microwells are designed with ultra-thin (∼40 μm) bases (floors). With the vacuum manifold mated to the bottom of the multi-layer assembly, the PDMS microwell floor flexes outward upon application of suction and material is pulled – via the trapdoor – from the top Bis-gel microwell into the receiving PDMS microwell (**Figure 1A**).

**Figure 2.**
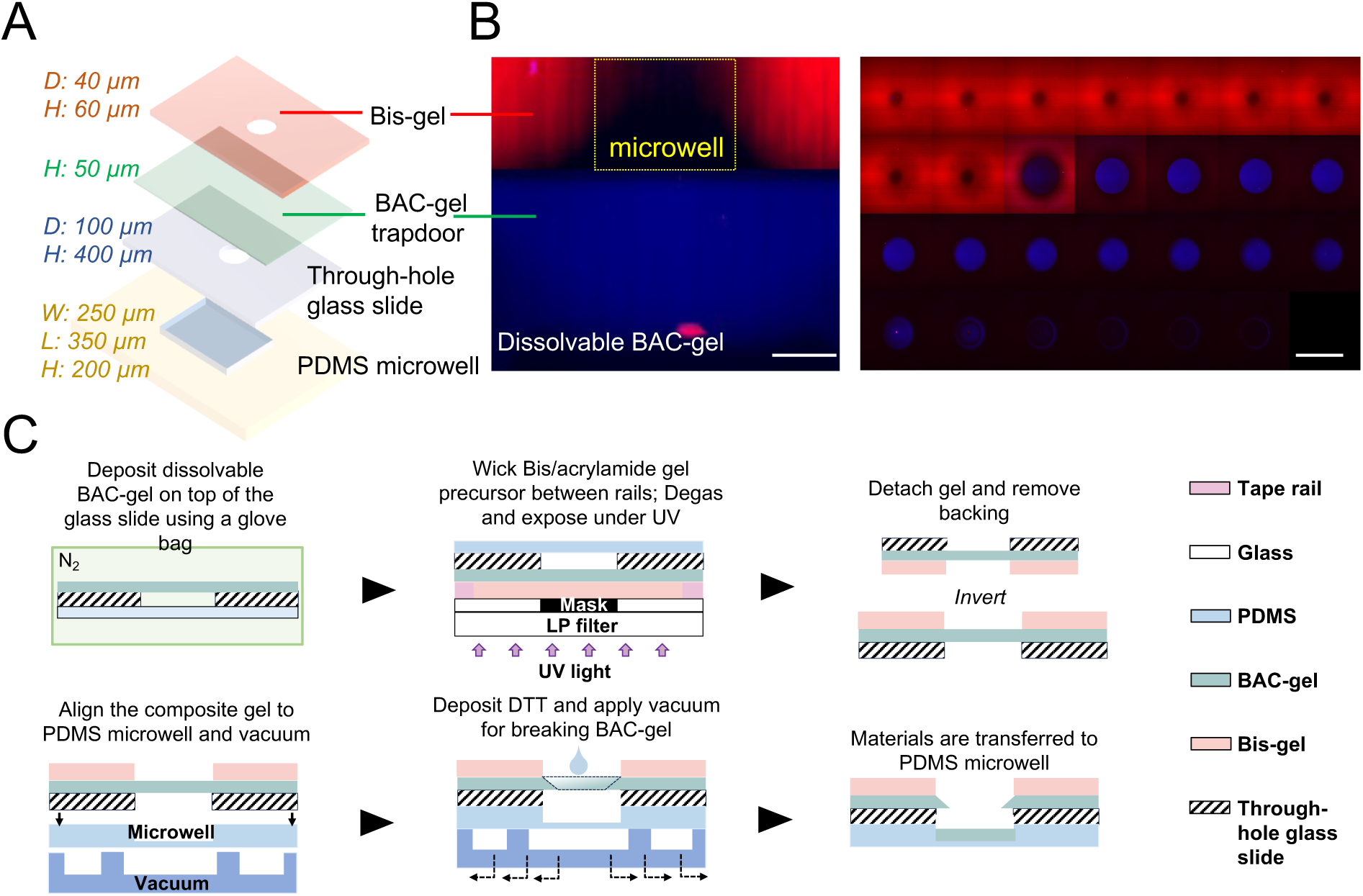
A multi-stage photopolymerization process fabricates a stacked microwell pair with trapdoor through connect. (**A**) Detailed schematic of each layer in VacTrap. Sequential layers include a 60 μm height and 40 μm diameter Bis-gel microwell (orange), a 50 μm height layer of dissolvable BAC-gel (green), 400 μm-thick through-holes with a 100-μm diameter through-hole glass slide, and a 350 by 250 μm receiving PDMS microwell. (**B**) Confocal imaging highlights the cross-sectional view (left, scale bar: 40 μm) and confocal sectioning of the Bis-gel with the BAC-gel trapdoor features (right, scale bar: 100 μm) with each z-section spanning 5 μm in the montage. Bis-gel microwell was labeled with Rhodamine B methacrylate (red) and the dissolvable BAC-gel was labeled with FITC Acrylate (blue). Images were taken with a 40x water-immersion Plan APO objective. **(C)** Schematic of the gel fabrication and dissolution process. The 50 μm height BAC-gel is chemically photopolymerized onto a through-hole glass slide within a nitrogen glove bag, followed by the fabrication of the Bis-gel atop the BAC-gel using photopolymerization employing a photomask and UV exposure. Gel dissolution is initiated by the application of DTT and vacuum suction force.

### Alignment strategy for fabrication of the multi-layered, interconnected VacTrap

For nucleus transfer to be successful, 40-μm diameter top-layer Bis-gel microwell must be polymerized and aligned atop of each 100-μm diameter glass through hole coated with 50 μm- thick layer of the BAC-gel that will function as a trapdoor conduit to the receiving 250 by 350-μm PDMS microwell (**Figure 2C**). The BAC-gel here will act as a temporary base of the Bis-gel microwell. Alignment must be achieved to sufficient precision across the 15 by 15 mm, 256 Bis-gel microwell array. One time-sensitive constraint arises: How to delay the polymerization and formation process of the Bis-gel microwell until their location is defined to align with each through-hole of the glass slide? To achieve this balance of timing, we implemented two distinct polymerization methods for BAC-gel and Bis-gel, each having a different time constant for polymerization. Since there is no restriction to the location or polymerization time of the BAC-gel, we employed the chemical polymerization for the BAC-gel layer. The BAC-gel was polymerized on top of the through-hole glass slide using acrylamide monomers and BAC as a cross-linker through free radical polymerization with tetramethylethylenediamine (TEMED) and Ammonium Persulfate (APS). To precisely position the Bis-gel microwells directly over the BAC-gel-coated through holes—ensuring the effective transfer of nuclei across multiple layers of the VacTrap system—the Bis-gel was photopolymerized using 2,2- Azobis[2-methyl-N-(2-hydroxyethyl)pro-pionamide] (VA-086, 1%) as a photoinitiator [40] A photomask was employed to define the microwells diameter and location. Under UV exposure, the Bis-gel precursor in the transparent regions of the mask was exposed and polymerized, while the opaque regions (containing 256 circular features, each 40 µm in diameter) blocked UV light, preventing polymerization and forming the microwells (**Figure 2C**). This photopolymerization approach allowed sufficient time to align the mask with the through holes, ensuring the Bis-gel microwells were accurately positioned over the BAC- gel-coated through holes, forming a composite gel. To initiate the transfer, the composite gel was then aligned to the PDMS microwell and the vacuum manifold using brightfield microscopy. Our vacuum manifold utilizes trapezoid pillars (surrounding each PDMS microwell) to prevent the PDMS microwell from collapsing when a vacuum force is applied (**Figure 1C**). This configuration ensures continued contact between the through-hole glass slide and the PDMS microwell.

### Design and fabrication of the trapdoor features

The diameter of the top-layer whole-cell receiving Bis-gel microwells is designed to closely match the diameter of individual mammalian cells (∼30-40 μm). To achieve an aspect ratio (1.3) designed to reduce the likelihood of capturing multiple mammalian cells in each microwell, we fabricate 60-um tall Bis-gel microwells [30].

To enhance the cell-settling efficiency, the Bis-gel whole-cell receiving layer is dehydrated prior to introducing a cell suspension. Drying polyacrylamide microwell results in the microwell diameter expanding upon dehydration by ∼1.5× for gels chemically polymerized (e.g., TEMED, APS). Deviations from an aspect ratio of ∼1.3 lead to ≥1 cell per microwell occupancy, which is not desired in single-cell resolution assays or sample preparation. Consequently, for a photopolymerization (versus chemical polymerization) process, we sought to understand the effect of UV dose (energy × exposure duration) on photopolymerization of the Bis-gel atop the dissolvable BAC-gel layer.

We asked what range of UV doses minimize Bis-gel expansion after dehydration, while preserving a target hydrated Bis-gel microwell diameter of 40 μm. All the while, the process maintains the Bis-gel layer as co-planar on top of the polymerized dissolvable BAC-gel in such a way that (1) the BAC-gel fully covers the top of the glass through holes and (2) the Bis-gel microwells are each aligned with the through holes in the glass slide (**Figure 3A**). Across a wide UV-dose range (1400 - 2000 mJ/cm²), we measured a ∼1.5× expansion in diameter for the Bis-gel microwells after dehydration when polymerizing using the lowest UV doses (1400 mJ/cm² and 1600 mJ/cm²) (**Figure 3B**). At 1400 mJ/cm², we observed darkening beneath the microwells by brightfield microscopy, particularly when approaching the through-hole glass slide during a z-axis sweep. We attributed the observation to potential under-polymerization of the Bis-gel, as indicated by a 57% increase in diameter after dehydration (0_hydrated_ = 37.2 ± 3.5 μm, 0_dehydrated_ = 58.6 ± 5.0 μm, N=100) (**Figure 3C**). In contrast, a higher UV dose of 2000 mJ/cm² resulted in a 25% shrinkage of the microwell diameter, with both hydrated and dehydrated microwells being too small for single-cell encapsulation (0_hydrated_ = 31.7 ± 3.1 μm, 0_dehydrated_ = 26.1 ± 2.8 μm, N=100). Optimal results were achieved at a UV dose of 1700 mJ/cm², where the diameter of 100 microwells was measured at 0_hydrated_ = 35.3 ± 1.8 μm and 0_dehydrated_ = 46.1 ± 3.3 μm, thus maintaining the target microwell diameter of ∼32 and 40 μm before and after drying, respectively, as is suitable for single mammalian cell encapsulation.

**Figure 3.**
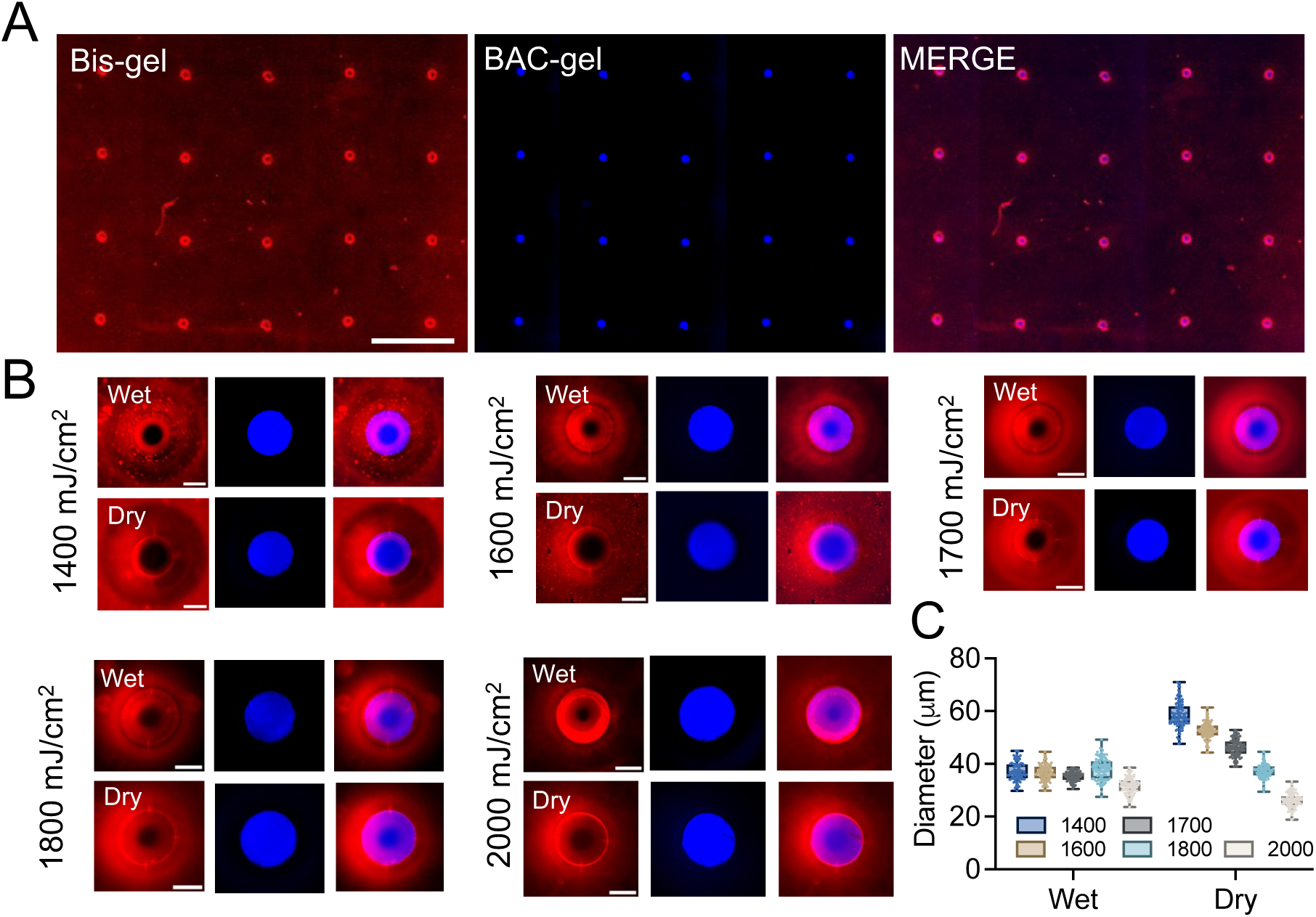
Control of UV doses is essential to fabricating the whole-cell receiving Bis-gel microwell array layer aligned with trapdoor features in an aligned BAC-gel layer which is seated on a glass support engraved with through holes. For visualization, Bis-gel was co-polymerized with 0.2 mM Rhodamine B methacrylate while BAC-gel was co-polymerized with 0.2 mM fluorescein-o-acrylate (FITC acrylate). (**A**) Fluorescence micrographs of the whole-cell receiving Bis-gel microwell array layer (red) and BAC-gel-on-glass trapdoor features (blue), along with merged image. Scale bar: 1 mm. (**B**) Characterization of hydrated and dehydrated Bis-gel microwell features photopolymerized under a range of 360-nm UV doses. Dehydration results in a slight expansion of features, as expected. Scale bar: 50 μm. (**C**) Diameter of resultant Bis-gel microwells fabricated using a range of UV doses, for hydrated and dehydrated imaging conditions (N=100 microwells for each condition). The diameter of the hydrated and dehydrated microwells were closest to the desired 40 μm diameter at a UV dose of 1700 mJ/cm², for this formulation.

Based on these observations, we posit that increasing the UV dose increases the Bis-gel stiffness and, thus, reduces the susceptibility of Bis-gel microwells to expansion upon dehydration. Previous research by Sheth et al [41] determined that the Young’s modulus – a measure of the stiffness of a hydrogel – is directly proportional to UV dose. The study considered photopolymerization of polyacrylamide hydrogels with the photoinitator Irgacure 2959 across a UV-dose range of 1500-2600 mJ/cm². The proposed underlying mechanism implicates higher UV dose to enhanced crosslinking reactions, resulting in formation of more functional crosslinks in the resultant gel versus those observed in a lower UV dose process. The crosslinks increase gel stiffness and, for our purposes, make the hydrogel less likely to shrink upon dehydration.

### Chemico-mechanical actuation of trapdoors to open fluidic connection between stacked microwell layers

We next sought to identify chemical and mechanical conditions well suited to actuating physical transfer of a nucleus through each trapdoor feature. Previous research has reported 10-20 mM DTT dissolving 0.392-0.500% BAC- gels in 1-5 min [39, 42]. However, these previous studies have considered dissolution of BAC-gel in a bulk form or as BAC-gel droplets immersed in a DTT solution, with thermoshaking. [39, 42]. Our layered microfluidic device presents a materially different dissolution environment for DTT-actuated BAC-gel dissolution. In our layered system, DTT must diffuse from point of application, through a 60-μm deep Bis-gel microwell, and then dissolve the 50-μm thick BAC-gel layer from the top.

In tandem with considerations of the BAC-gel composition, we considered compatible approaches to apply force to the dissolving BAC-gel and expedite formation of a fluidic interconnection between the two layers. Primary among our considerations was a vacuum-driven force wherein we attached a microfluidic vacuum manifold underneath the nucleus-receiving PDMS microwell layer. PDMS casting fabricated a thin PDMS floor in each nuclei-receiving PDMS microwell (thickness∼40 μm). The thin PDMS floor is important to provide physical compliance sufficient to effectively transfer vacuum- generated suction force from the vacuum manifold layer to the contents of the PDMS microwell, the BAC-gel in the glass through holes, and finally into the upper Bis-gel microwell compartment. The vacuum manifold generates continuous negative pressure across the gas-permeable PDMS thin floor, allowing air to diffuse from the Bis-gel microwell through the BAC-gel and glass through-holes, which in turn drives DTT flow through the Bis-gel microwell, efficiently dissolving the BAC-gel.

To prevent collapse of the PDMS microwell and floor when vacuum force is applied, we employed an array of structural-support pillars surrounding each PDMS microwell, thereby ensuring supportive structural contact between the through-hole glass slide and the PDMS microwell layer (**Figure 4A**). We first employed circular cross-section pillars and observed physical behavior and features when the BAC-gel layer incorporated a fluorescently labeled acrylamide monomer. However, we found that the circular cross-section pillars did not prevent PDMS microwell deformation under application of the vacuum force (**Figure 4A**). Circular cross-section pillars resulted in detachment between the PDMS microwell and the vacuum manifold. In contrast, when the larger-surface area trapezoidal cross-section pillars were employed, the PDMS microwell was not observed to either deform or detach, and fluidic connectivity was observed between the stacked Bis-gel and PDMS microwells. Consequently, we opted to utilize trapezoidal cross-section support pillars around the PDMS microwells.

**Figure 4.**
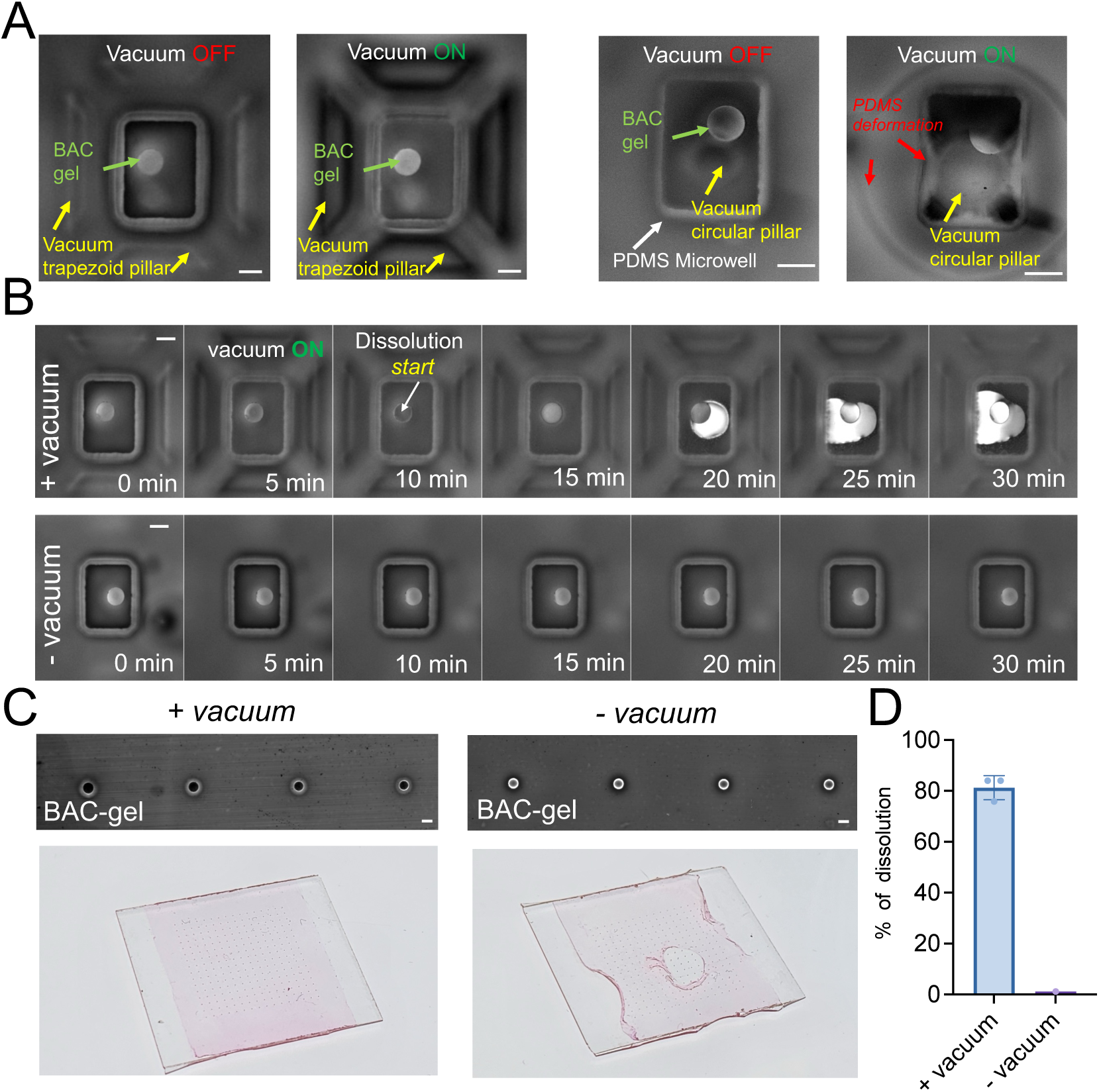
Chemico-mechanical actuation opens the cell-scale trapdoor feature for the physical transfer of single nuclei from the top microwell to the bottom microwell in a VacTrap microwell stack. (**A**) Fluorescence micrographs of pre- and post-actuation of the trapdoor feature by an applied suction force levied by the vacuum manifold with trapezoid and circular pillars. For visualization, the BAC-gel is copolymerized with 0.2 mM FITC acrylate in all fluorescence images reported in this Figure. (Left) Fluorescence micrographs show deformation and detachment of circular structural pillars after suction is applied to the PDMS microwell by the vacuum manifold. (Right). Fluorescence micrographs show the structural integrity of trapezoidal pillars and PDMS microwell after suction is applied to the PDMS microwell by the vacuum manifold. Scale bar: 100 µm. (**B**) Timelapse of fluorescence micrographs of the PDMS microwell and trapdoor feature report dissolution of the 0.2% BAC-gel trapdoor with application of 40 mM DTT, observed with (+ vacuum) and without (- vacuum) vacuum application. Fluorescence microscopy uses a FITC (488 nm) filter focusing on BAC-gel trapdoor. Scale bar: 100 µm. (**C**) BAC-gel trapdoor dissolution efficacy with and without vacuum applied (+vacuum, -vacuum). Micrograph imaging (Scale bar: 100 µm) using Genepix microarray scanner shows the complete and specific dissolution of BAC-gel around the through-hole area, indicated by the loss of fluorescence in the through-hole area. Without the vacuum, the dissolution is limited, and the fluorescence signal of the BAC-gel remained around the through-hole. Moreover, the non-specific dissolution of the BAC-gel causes gel damage and detachment due to loss of support beneath the Bis-gel. (**D**) BAC-gel dissolution efficiency as determined by enumerating PDMS microwells exhibiting circumscribed FITC signal, indicative of successful BAC-gel dissolution and physical transfer into the nuclei-receiving PDMS microwell.

To understand the importance of not just dissolving the BAC-gel comprising the trapdoor, but also applying a gentle suction force on that depolymerized BAG-gel bolus, we conserved the dissolution process with and without applied vacuum from the vacuum manifold (**Figure 4B**) for a BAC-gel trapdoor fabricated with 0.2% BAC crosslinker, using 40 mM DTT. With the vacuum applied, dissolution through a 50-μm-thick BAC-gel layer occurred within 10 min (**Figure 4B**). Upon activation of the vacuum, the DTT rapidly reached the microwell, indicated by the reduction in fluorescence signal from the BAC- gel on top of the through-hole glass. Within the subsequent 5 min, dissolution commenced. Dissolution was considered complete when the fluorescence signal from the BAC-gel reached its maximum before a reduction in fluorescence. The signal trend indicates that the dissolved fluorescent BAC-gel initially accumulates at the glass through holes, resulting in a peak fluorescence signal. As the gel continues to dissolve, the liquified gel passes through the through-hole and into the underlying PDMS microwell, causing a decrease in fluorescence as the material moves out of the focal plane. With the vacuum support, the BAC-gel around the through holes is fully dissolved, indicated by the loss of fluorescence in the through-hole area (**Figure 4C**). In contrast, when a trapdoor feature composed of 0.2% BAC was exposed to 40 mM DTT without application of an external mechanical force (-vacuum), the BAC-gel layer did not dissolve and transfer into the PDMS microwell, and no fluidic nor materials connection was observed between the stacked Bis-gel and PDMS microwells after 30 min (**Figure 4B**), as evidenced by the fluorescence signal of BAC-gel remaining around the through-hole (**Figure 4C**). The vacuum facilitated dissolution in an average of 80% of microwells (∼179 microwells). Without applied vacuum, dissolution of the BAC-gel was observed in less than 1% of microwells (**Figure 4D**). Dissolution efficiency depends on the precise alignment of the Bis-gel microwell and trapdoor feature with the lower-layer PDMS microwell to ensure the suction force is transmitted effectively through the microwell stack. Ensuring timely dissolution of the BAC-gel is crucial to maintaining the integrity of the Bis-gel microwell. Without specific dissolution within the through-hole area only, the Bis-gel can detach due to a loss of structural support from the BAC-gel and the through-hole glass slide (**Figure 4C).** These observations suggest the significance of applying suction to facilitate fluidic interconnection between the stacked microwell layers.

To understand the practical implications of dissolving a BAC-gel in a layered configuration, we studied parameters that influence the dissolution rate, including BAC concentration, DTT concentration, and UV dose used in Bis-gel photopolymerization (**Figure 5**). We first tested a range of BAC crosslinker concentrations from 0.150% to 0.400%. Higher BAC concentrations resulted in a stiffer gel exhibiting a longer time to dissolve. Therefore, we aimed for a low BAC concentration to facilitate rapid dissolution of the BAC-gel layer while still maintaining the integrity of the Bis-gel microwell and dissolvable trapdoor feature. We observed that a 50-μm thick BAC-gel with 0.150% BAC can be dissolved by 100 mM DTT in 3 min (**Figure 5A**), which was in the target dissolution-performance range. However, this lower BAC concentration made the gel more susceptible to tearing during the fabrication process. With that in mind, a 0.200% BAC was observed to dissolve within 3- 5 min using 100 mM DTT (**Figure 5A**), still within the desired timeframe but with enhanced mechanical robustness which is helpful for reliable fabrication. Taken together, a 0.2% BAC-gel was selected for further analysis.

**Figure 5.**
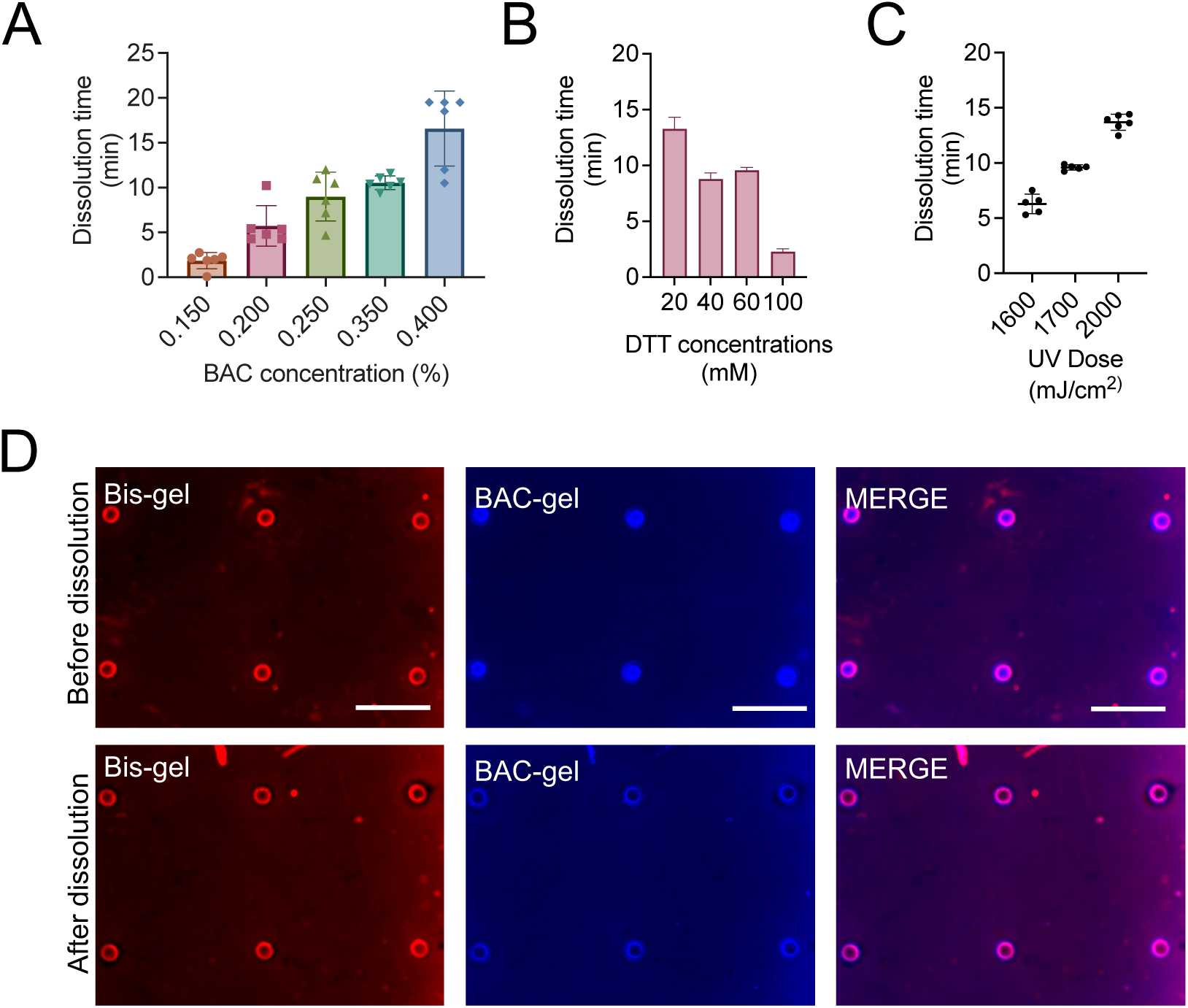
Chemico-mechanical actuation of the BAC-gel trapdoor is achieved within 3-5 min with 0.2% BAC-gel and 100 mM DTT, ensuring rapid fluidic connectivity and materials transfer between the stacked microwells. (**A**) Dissolution time of the BAC-gel trapdoor as a function of BAC concentration with [DTT] = 100 mM. (**B**) Dissolution time of the BAC-gel trapdoor (0.2% BAC) as a function of applied DTT concentration. (**C**) Dissolution time of the 0.2% BAC-gel trapdoor with 40 mM DTT as a function of UV dose used for Bis-gel microwell photopolymerization. Fixed acrylamide concentration of 6% w/v. (**D**) Fluorescence micrographs of the Bis-gel with microwells stained with 0.2 mM Rhodamine B methacrylate, BAC-gel layer stained with 0.2 mM fluorescein acrylate, and merged images after BAC-gel dissolution with 100 mM DTT and an applied vacuum force did not affect the integrity of the Bis-gel microwell. Scale bar: 500 µm.

In tandem, we considered a range of DTT concentrations from 20 mM to 100 mM for dissolution of trapdoor features fabricated with 0.2% BAC-gel, with complete dissolution achieved in 3 min with 100 mM DTT. Application of 20 mM of DTT required nearly 15 min for dissolution (**Figure 5B**). However, DTT is a common redox reagent used to break down protein disulfide bonds, including antibodies [43]. Therefore, for potential proteomics applications in the PAG microwell layer, we sought to reduce DTT concentrations to 40 mM, followed by several washes with high pH buffer (>8) at high temperature to deactivate DTT before immunoprobing. DTT does not interfere with PCR or reverse transcription, making this dissolvable gel suitable for common genomic and nucleic acids applications such as DNA or RNA-seq [39].

Surprisingly, we found that the UV dose used for Bis-gel photopolymerization did affect the dissolution of the underlying BAC-gel trapdoor, with increasing UV dose increasing the required dissolution time (**Figure 5C**). However, choosing a low dose of UV for Bis-gel photopolymerization could lead to underexposure causing microwell expansion and incomplete polymerization beneath the Bis-gel microwell (**Figure 3B**). We hypothesize that UV-based activation homolytically cleaves disulfide bonds to yield two separated thiol radicals [44, 45]. While disulfide bonds could reform if the radical species generated remain in proximity after cleavage, the radicals may recombine with different thiol radicals within the gel matrix, not necessarily from the same original disulfide bridge. Such recombination would cause an observed temporal delay in dissolution. Moreover, excess photoinitiator (VA-86) trapped in the Bis-gel may lead to further polymerization of the BAC- gel around the microwell area, causing further delay in dissolution. With 100 mM of DTT, a 0.2% BAC concentration, and a 1700 mJ/cm^2^ UV exposure for the Bis-gel, dissolution was completed in < 5 min without any detectable damage to the Bis-gel microwell after dissolution (**Figure 5D**).

### Actuation of trapdoor features allows concurrent physical transfer of isolated nuclei

We sought to understand the capability of VacTrap to transfer isolated nuclei through the dissolved BAC-gel while maintaining the physical integrity of the nucleus after an applied (suction) mechanical force. To assess simple physical integrity of isolated nuclei, we employed fluorescence microscopy to inspect whether transferred nuclei were physically intact or physically compromised after transfer through a 0.2% BAC-gel trapdoor dissolved by applying 100 mM DTT and suction. **Figure 6A** illustrates nucleus transfer through the trapdoor of the BAC-gel into the nuclei-receiving PDMS microwell. Nuclei were observed transferring into the nucleus-receiving PDMS microwells at ∼360 s after vacuum activation while the dissolution began first at ∼135 s. By fluorescently labeling both the BAC-gel in the trapdoor feature and the isolated nuclei with HOECHST 33342 we observed nuclei transferred in nearly 80% of PDMS microwells inspected (**Figure 6B and C**).

**Figure 6.**
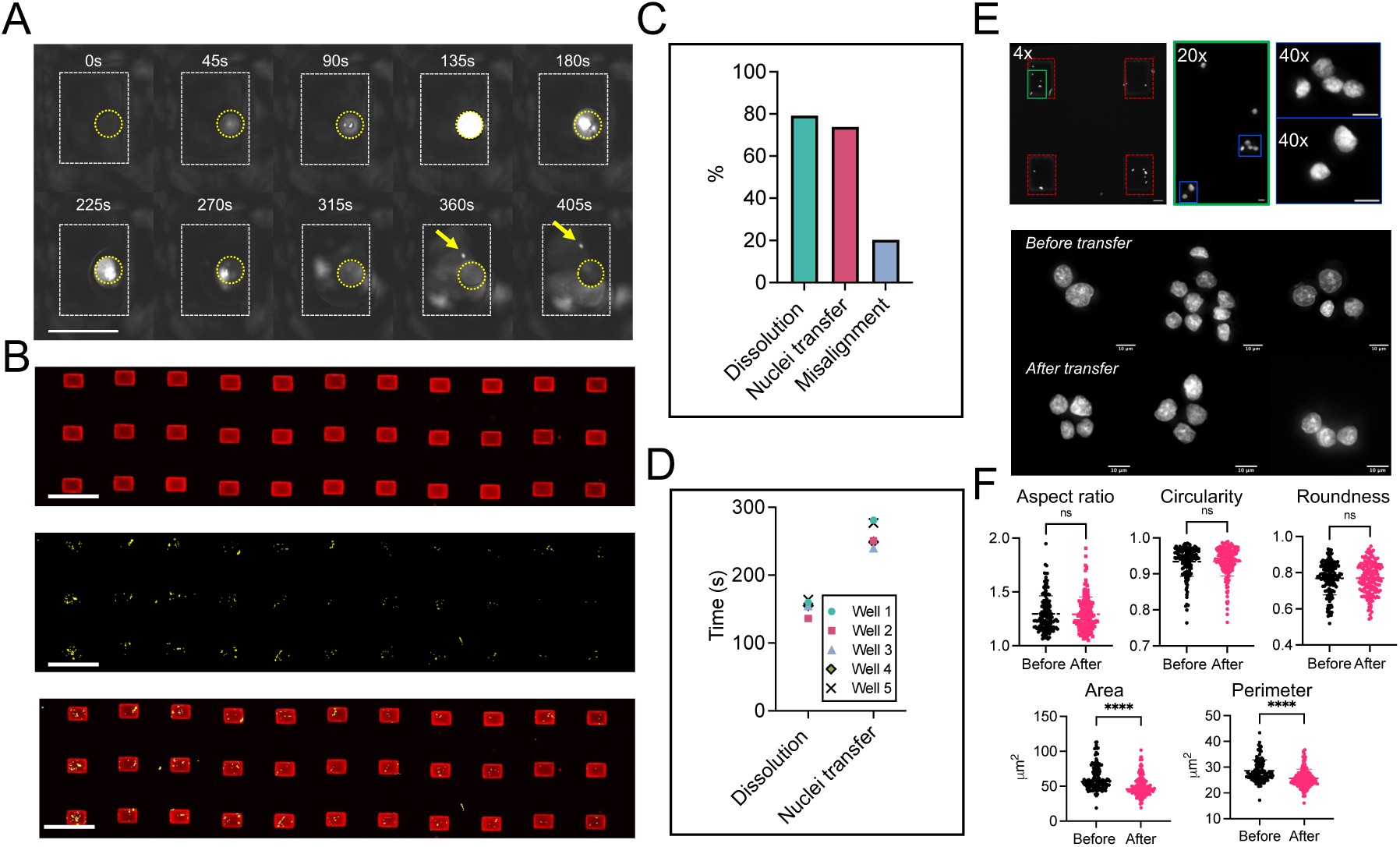
VacTrap simultaneously transfers single mammalian nuclei across an array of stacked microwells while maintaining nuclear morphological integrity. (A) Timelapse of fluorescence images showing transfer of isolated nuclei from breast cancer cell line cells (MCF7). For visualization, nuclei were labeled with 40 µM HOESCHT 33342, settled into the cell-receiving Bis-gel microwells. FITC-acrylate labeled BAC-gel trapdoor was activated with 100 mM DTT followed by vacuum activation. Timelapse imaging was conducted using a 4x PLAN APO objective in 1 second intervals. Imaging utilized a UV filter set (365 nm) focused on the labeled nuclei. Scale bar: 250 µm. (B) False-color fluorescence micrographs of the receiving PDMS microwell after nuclei transfer (yellow) concurrent with trapdoor BAC-gel dissolution (red). Scale bar: 1 mm. From top to bottom: Dissolved trapdoor of BAC-gel (red), transferred nuclei (yellow), and merged micrographs. (C) Microscopy-based analysis of nucleus transfer yield shows the percentage of microwells (%) showing both trapdoor dissolution and successful nuclei transfer are nearly identical, with discrepancies attributed to misalignment between the nucleus-receiving PDMS microwell on the bottom layer and the whole-cell receiving Bis-gel microwells on the top device layer. (D) Dissolution time and nuclei transfer across 6 representative trapdoor features suggests nearly synchronized trapdoor actuation across the microwell array. (E) Fluorescence microscopy inspection of transferred nuclei housed in the nucleus-receiving PDMS microwell array suggests nuclei remain intact after transfer. Scale bar: 100 µm (4x) and 10 µm (20x and 40x). (F) Morphological analysis of transferred nuclei before and after trapdoor-assisted transfer that relies on a combination of trapdoor dissolution (DTT) and applied suction using the integrated vacuum manifold layer.

To understand the degree of synchronization in the dissolution times across the trapdoor features in a microwell array (**Figure 6D**), we monitored dissolution with fluorescence microscopy and measured trapdoor BAC-gel dissolution times ranging from 136-160 s, with an average of 154 +/- 10 seconds (N=6) with nucleus transfer occurring at ∼105 +/-15 seconds post-dissolution. The observed delay between the initial dissolution of the BAC-gel trapdoor and nucleus transfer arises from the requirement for complete dissolution of the BAC-gel, which requires 3-5 min with 100 mM DTT. Ensuring simultaneous dissolution and transfer is essential for maintaining the integrity of the nuclei throughout the entire microwell array.

To assess overall yield of trapdoors with suitable performance, inspection of the microwells by microscopy during BAC-gel dissolution revealed that ∼80% of the BAC-gel microwells dissolved, corresponding with the percentage of nuclei transferred within the same microwell array (**Figure 6C**). The high but not perfect yield in functional trapdoor features is attributed to misalignment between the stacked pair of Bis-gel and PDMS microwell arrays. Additionally, PDMS is known to shrink when cured at high temperatures such as those used in this study, so curing PDMS microwells at room temperature should reduce shrinkage and enhance alignment accuracy.

### Transfer of isolated nuclei from single mammalian cells

Finally, to extend understanding beyond the physical integrity of extracted nuclei, we sought to assess nuclear phenotype (e.g., morphology). Here, we leveraged the transparency of the PDMS microwells to assess the morphology of nuclei before and after transfer (**Figure 6E-F**). Common morphological parameters including nuclear aspect ratio, circularity, roughness, area, and perimeter were analyzed (**Figure 6F)**. Our results indicate no detectable changes in aspect ratio, circularity, or roughness before and after nuclei transfer. However, alterations in area and perimeter were observed. We hypothesize that the changes in area and perimeter are attributable to the response of nuclei upon exposure to DTT during the transfer process as well as imaging artifacts that arise from imaging through the PDMS microwells. Nevertheless, nuclei remained intact and retained overall shape.

## Conclusions

In this study, we introduced the VacTrap system, a multilayer microfluidic device designed to facilitate high throughput, spatially indexed transfer of nuclei from each cell of hundreds of single cells. By integrating a stacked pair of microwells – a top layer that is the cell-receiving Bis-gel microwell array and a bottom layer that is the nucleus-receiving PDMS microwell array – with interleaving dissolvable trapdoor features and a vacuum-driven force actuation system, VacTrap simultaneously extracts and transfers isolated nuclei across hundreds of microwells within 3-5 min. Importantly, VacTrap preserves nuclear integrity and indexing each nucleus back to its originating cell and any associated data collected on that intact cell prior to nuclear extraction.

As a sample preparation device, VacTrap automates the functions of isolating and measuring (e.g., imaging) individual intact cells, fraction of the nucleus from each imaged cell, and then synchronized physical transfer of the isolated nuclei into a PDMS microwell array suitable for subsequent nuclei measurement and analysis (e.g., PCR). To provide a compartment for intact cell imaging and nuclei fractionation, and one for harsh chemical manipulation of isolated nuclei, VacTrap is designed with a stacked microwell array comprising: 1) a top-layer of Bis-gel microwells used to isolate the originating single cell and 2) a bottom-layer of PDMS microwells used to compartmentalize each extracted nucleus. Once the nucleus has been fractionated from the originating cell, the VacTrap system establishes a fluidic connection on demand between the stacked microwell pair using chemical and mechanical actuation. Spatial indexing of hundreds of originating intact cells to their resultant fractionated nuclei takes advantage of the microarray layout, which is compatible with time-lapse imaging. A precision single-cell preparation technique, VacTrap enhances the throughput of organelle isolation (here, demonstrated for nuclei) and ensures the rapid and reliable transfer necessary for downstream multi omics analyses, thus demonstrating potential for inclusion in single-cell multi omics research tools.

## Experimental Section

### Chemicals

Tetramethylethylenediamine (TEMED, T9281), 40% Acrylamide solution (A4058), Acrylamide/Bis-acrylamide 40% solution (29:1), N,N’-Bis(acryloyl)cystamine (A4929), Ammonium persulfate (APS, A3678), Methanol (34860), Dimethylsulfoxide (DMSO, D2438), Fluoresceine O Acrylate (568856), and 3-(Trimethoxysily)propyl methacrylate were obtained from Sigma Aldrich.

0.1M Dithioerythritol (DTT) solution, Phosphate buffered saline (PBS, 10010023), and Hoechst 33342 solution (20 mM, 62249) were purchased from Thermo Fisher. BP-APMA (BPMAC) was custom-synthesized by Raybow. Photoinitiator 2,2-Azobis(2-methyl-N-(2- hydroxyethyl)propionamide) (VA-086) was acquired from FujiFilm Wako Pure Chemical Corporation. Gel Slick was purchased from Lonza (#50640). Molecular biology grade water was from Corning (46-000-CV). Tris-glycine (10×) buffer (25 mM Tris, pH 8.3; 192 mM glycine) was obtained from Bio-Rad (#1610734). Methacryloxyethyl thiocarbamoyl rhodamine B (Rhodamine B methacrylate, 23591-100) was purchased from Polysciences. IGEPAL® CA-630 MegaPure™ Detergent, 10% solution was acquired from Abcam (ab285400). 10% Tween-20 Dnase/Rnase Tested, Sterile was from Teknova (Teknova T0027). Digitonin solution supplied at 20 mg/ml in DMSO was purchased from Promega (G9441).

### Buffers

Nuclei were isolated using ATAC-RSB buffer [46] which was prepared by mixing 500 µL of 1M Tris-HCl pH 7.4 (10 mM), 100 µL of 5M NaCl (10 mM), 150 µL of 1M MgCl_2_ (3 mM), and 49.25 mL of molecular biology grade water. The lysis buffer was prepared by adding 0.1% IGEPAL CA-630, 0.1% Tween-20, and 0.01% digitonin to ATAC-RSB buffer to reach the final volume. The wash buffer contained 0.1% Tween-20 in ATAC-RSB buffer.

### Fabrication of the trapdoor BAC-gel layer on through-hole glass slides

100-μm diameter, 400-μm thick through-hole glass slides (28 mm by 40 mm) were generously provided by Arralyze (LPKF Laser & Electronics AG, Germany). BOROFLOAT® 33 (Schott AG), a borosilicate glass that is widely used for life science applications due to low autofluorescence and high optical transparency in the visible region, was used as a substrate. Details on the Laser Induced Deep Etching protocol can be found in previous studies [36]. The slides were silanized to enhance the gel bonding on the glass slide by adding a methacrylate group as previously described [30].

BAC-gel fabrication was performed in a glove bag (Thermo Scientific, 093737.LK) with continuous nitrogen flow to prevent oxygen inhibition of polymerization. 50-μm Kapton tape rails (315-CQT-0.250-ND) were taped onto a large glass slide (Ted Pella, 260234- 25) with 20 mm spacing to align the through-hole area. The glass slide was washed with IPA and dried with nitrogen. Gel slick^R^ (Lonza) (600 μL) was spread between the Kapton tape rails and dried at room temperature. The glass slide was then washed with water and dried with a Kimwipe using a buffing motion to remove excess gel slick.

A 20 mm x 18 mm PDMS membrane was cut and applied to one side of the through-hole glass slide to limit gel precursor diffusion during BAC-gel fabrication. The through-hole glass slide was taped atop the Kapton tape rails on the large glass slide with the PDMS membrane facing up. At least ∼5 mm from each long side of the through-hole glass slide should sit on top of the Kapton tape rails, resulting in the gel-free edges after fabrication. The assembly was degassed for 10 min before being moved to a glove bag until the gel precursor was ready.

10% APS (w/v) and 10% TEMED (v/v) were prepared with molecular biology grade water and moved to the glove bag. BAC solution was made by dissolving ∼22 mg of BAC in 100% methanol, followed by vortexing. BAC-gel precursor was prepared with 6% (w/v) acrylamide, 1× Tris-Glycine (pH 8.3), molecular grade water, and various concentrations of BAC (0.150%, 0.200%, 0.250%, 0.350%, and 0.400%). For some experiments, 100 mM fluorescein o-acrylate in DMSO was added to the gel precursor for a final concentration of 0.2 mM. The gel precursor was degassed and sonicated for 15 min before adding 10% APS and 10% TEMED at a final concentration of 0.1% under the glove bag. 1 mL of gel precursor was quickly wicked through the through-hole glass slide and polymerized under nitrogen for 20 min. After polymerization, the through-hole glass slides and the PDMS membrane were removed, and the gel was incubated with DI water for 5 min before removal from the rails. The gels were kept in water for at least 2 hours before use.

### Fabrication of the Bis-gel microwells with photopolymerization

A customize 8×8 photomask (Artnet Pro) with 40-μm-diameter dark circular features and transparent fields was affixed to heat-resistant borosilicate glass (8" × 6", 1/8" thickness, McMaster Carr 8476K72). Two pieces of 60-μm-thick Kapton tape (3M 5419) were applied to form two rails for Bis-gel fabrication, set to cover the through-holes and BAC-gel area.

Gel slick (400 μL) was applied between the rails and dried at room temperature for 3 min. Excess gel slick on the mask was cleaned with a Kimwipe using a circular buffing motion. On the other side of the glass plate, a long pass filter sheet (8" × 6") was cut and fixed with Kapton tape.

The BAC-gel was dried with nitrogen before attaching a 20 mm × 18 mm PDMS membrane to cover the through-hole. Two pieces of 50-μm Kapton tape were applied to the gel-free edges (∼5 mm wide) of the through-hole glass slide and cut to shape. These rails compensate for BAC-gel height expansion when exposed to the Bis-gel precursor. The entire assembly was moved to a vacuum chamber and kept closed without vacuum until the gel precursor was ready.

VA-086 photoinitiator was dissolved in water to a final concentration of 2% (w/v). Bis-gel precursor was prepared with molecular grade water, 7% Acrylamide/Bis-acrylamide (29:1), 3 mM BPMAC in DMSO, 1× Tris-glycine (pH 8.3), and 1% VA-086, adjusted with molecular biology grade water. To stain the gel, 100 mM Rhodamine B methacrylate was added to the precursor for a final concentration of 0.2 mM. The gel precursor was degassed for 10 min before wicking through the BAC-gel through-hole glass slide and vacuuming until all bubbles were removed.

The glass plate was then placed under an OAI UV exposure system (Optical Associates, Incorporated) with UV power of ∼20 mW/cm² (OAI UV Probe 365nm, measured without the long-pass filter) for doses of 1400, 1600, 1700, 1800, and 2000 mJ/cm². The standard dose for most experiments was 1700 mJ/cm² (∼85 s exposure with ∼20 mW/cm² UV energy). After photopolymerization, gels were incubated with water for 5 min before detachment. The composite gels were kept in molecular biology grade water until use.

### Soft lithography for fabrication of PDMS layers

The PDMS microwell array consists of 16 rows by 16 columns of rectangular microwells, each measuring 350 µm by 250 µm, with a 1 mm spacing center-to-center. The microwell SU-8 mastermold was fabricated using SU-8 2100 (Kayaku Advanced Materials) to achieve a height of 200 µm, following the manufacturer’s instructions. Then the PDMS microwells were produced by spinning approximately 5g of a 10:1 PDMS mixture on the SU8 mastermold in two steps: the first step for 5 seconds at 100 rpm with an acceleration time of 5 seconds, and the second step for 30 seconds at 400 rpm with an acceleration of 100 rpm, followed by 3 hours curing at 80 °C. Before any experiment, the PDMS microwell was deposited in the air plasma cleaner (PDC-32G, Harris plasma) with a vacuum setup of 0.470 torr using an ICME vacuum pump. The radiofrequency power was set to High for 3 minutes.

A vacuum manifold was prepared by casting approximately 27g of a 10:1 PDMS mixture onto a 100 µm height SU-8 mold using SU-8 3050 (Kayaku Advanced Materials), also according to the manufacturer’s instructions. The vacuum manifold featured trapezoid structures with bases of 250 µm and 443 µm, and legs of 147 µm. Prior to PDMS casting, the PDMS mixture was degassed for 1 hour before being applied to the wafers. All PDMS was cured at 80°C for 3 hours and allowed to cool to room temperature before use. The vacuum manifold had four outlets, which were created using a 2.5 mm biopsy punch (Integra) and connected to soft PVC Plastic Tubing for Air and Water, 1/32" ID, 3/32" OD (McMaster-Carr) for vacuum application.

### Nuclear isolation

Cancer cell line MCF7 Tet-off parental cells were kindly gifted by the Arribas Lab from the Vall’ d’Hebron Institute of Oncology. The cell line was authenticated by short tandem repeat profiling by the UC Berkeley Cell Culture facility and tested negative for mycoplasma. For each experiment, cell culture was maintained at 37 °C and 5% CO2 in Dulbecco’s Modified Eagle Medium/Nutrient Mixture F-12 (Gibco™ DMEM/F- 12, GlutaMAX™ supplement, Thermofisher, 10565018) supplied with 10% fetal bovine serum (Gemini Bio), 0.2 mg/ml Gibco™ Geneticin™ Selective Antibiotic (G418 Sulfate), and 1 μg/ml doxycycline (Sigma) until 80% confluency and detached with 0.05% Trypsin- EDTA (Gibco #25300-054) for 4-5 min.

1 million viable cells were aliquoted into 1.5 ml LoBind Eppendorf tubes. The cells were then centrifuged at 500 g for 5 min at 4°C. After centrifugation, the medium was removed, and the cells were resuspended in 1 ml of cold 1x PBS buffer. The cells were centrifuged again at 500 g for 5 min at 4°C, and the PBS was aspirated. Subsequently, 300 µL of cold lysis buffer was added to the sample, and the cells were mixed 10 times. The sample was incubated on ice for 5 min. After incubation, 1 ml of cold wash buffer was added to each sample, and the tubes were inverted 5 times to mix. The nuclei were pelleted with the hinge facing in at 500 g for 3 min at 4°C, then centrifuged again with the hinge facing out at 500 g for 3 min at 4°C. The supernatant was aspirated in two steps: 1000 µL was removed with a P1000 pipette, and the remaining 50-100 µL was removed with a P200 pipette. The nuclei were gently resuspended in 250 µL of wash buffer using a wide-bore tip (Rainin). The quality and count of the nuclei were assessed using a Countess (10 µL of nuclei with 10 µL of Trypan blue) To fluorescently label nuclei for imaging, 2 µL of 20 mM Hoechst was added into 1000 µL of PBS to prepare the staining wash buffer. An aliquot of 100,000 nuclei was added to 1000 µL of staining wash buffer and incubated for 20 min on ice. The nuclei were pelleted with the hinge facing in at 500 g for 5 min at 4°C, then centrifuged again with the hinge facing out at 500 g for 5 min at 4°C. The supernatant was aspirated, and the nuclei were resuspended in 1000 µL of PBS to achieve a concentration of approximately 100 nuclei/µL.

### Alignment and assembly of the VacTrap device layers

Before aligning the device, the composite gel was gently dried using nitrogen. The back of the glass slide was then cleaned with Scotch tape to ensure a seamless contact between the PDMS microwell and the composite gel. The composite gel was initially aligned with the PDMS microwell, then inverted, and the vacuum manifold was carefully applied to the back of the PDMS microwell.

### Actuation of the Bis-gel trapdoor features

The four outlets of the vacuum manifold were connected to tubing and a house vacuum system. Subsequently, 300 µL of DTT was delicately applied to the surface of the gel, followed by a 2-minute incubation period before activating the vacuum.

To assess nuclei transfer between the composite gel and PDMS microwell arrays, the nuclei were allowed to gently settle onto the gel for 10-15 min. Afterward, the gel was washed with PBS to remove excess nuclei, followed by the application of DTT and the activation of the vacuum.

### Imaging

All imaging reported in this study was performed using an Olympus IX51 microscope with various Plan Apo objectives (4x, 10x, 20x, and 40x) and filter sets for GFP, Texas Red, and UV (DAPI). Confocal imaging was conducted with a Bruker Confocal Microscope at the UC Berkeley QB3 Cell and Tissue Analysis Facility, utilizing an Olympus Plan APO 40x water immersion objective. Additionally, some imaging in this study was performed using a Genepix MicroArray Scanner (Genepix 4300A, Molecular Devices). Image processing was done using ImageJ, and nuclei morphology was analyzed with the MicrobeJ plugin.

## Data availability statement

The data that support the findings of this study are available from the corresponding author upon reasonable request.

## Acknowledgements

This work was funded by the National Institutes of Health (NIH) R01CA20301 (A.E.H.) and the Chan-Zuckerberg Biohub Investigator Award (A.E.H.). We appreciate the generous support from Arralyze company through the donation of through-hole glass slides for the initial concept development of VacTrap. We sincerely thank Paul Lum, Managing Director of the QB3 Berkeley Biomolecular Nanotechnology Center (BNC), and Dr. Mary West, Director of the Cell and Tissue Analysis Facility and the High Throughput Screening Facility, for their instrument training and supervision. We acknowledge all members of the Herr Lab at UC Berkeley.

## References

1. Liao, P.C., et al., Isolation of mitochondria from cells and tissues. Methods Cell Biol, 2020. 155: p. 3–31.

2. Drissi, R., M.-L. Dubois, and F.-M. Boisvert, Proteomics methods for subcellular proteome analysis. The FEBS Journal, 2013. 280(22): p. 5626–5634.

3. Andersen, J.S., et al., Directed proteomic analysis of the human nucleolus. Curr Biol, 2002. 12(1): p. 1–11.

4. Taylor, S.W., et al., Characterization of the human heart mitochondrial proteome. Nat Biotechnol, 2003. 21(3): p. 281–6.

5. Satori, C.P., V. Kostal, and E.A. Arriaga, Review on recent advances in the analysis of isolated organelles. Anal Chim Acta, 2012. 753: p. 8–18.

6. Parsons, H.T., S.M.G. Fernández-Niño, and J.L. Heazlewood, Separation of the Plant Golgi Apparatus and Endoplasmic Reticulum by Free-Flow Electrophoresis, in Plant Proteomics: Methods and Protocols, J.V. Jorrin-Novo, et al., Editors. 2014, Humana Press: Totowa, NJ. p. 527–539.

7. Wildgruber, R., et al., Free-flow electrophoresis in proteome sample preparation. PROTEOMICS, 2014. 14(4-5): p. 629–636.

8. Li, C.M., et al., Partial Purification of a Megadalton DNA Replication Complex by Free Flow Electrophoresis. PLoS One, 2016. 11(12): p. e0169259.

9. Ramsby, M.L., G.S. Makowski, and E.A. Khairallah, DiJerential detergent fractionation of isolated hepatocytes: biochemical, immunochemical and two-dimensional gel electrophoresis characterization of cytoskeletal and noncytoskeletal compartments. Electrophoresis, 1994. 15(2): p. 265–77.

10. Sawhney, S., R. Stubbs, and K. Hood, Reproducibility, sensitivity and compatibility of the ProteoExtract subcellular fractionation kit with saturation labeling of laser microdissected tissues. Proteomics, 2009. 9(16): p. 4087–92.

11. Hwang, B., J.H. Lee, and D. Bang, Single-cell RNA sequencing technologies and bioinformatics pipelines. Exp Mol Med, 2018. 50(8): p. 1–14.

12. Butto, T., et al., Nuclei on the Rise: When Nuclei-Based Methods Meet Next- Generation Sequencing. Cells, 2023. 12(7).

13. Sancho-Albero, M., et al., Isolation of exosomes from whole blood by a new microfluidic device: proof of concept application in the diagnosis and monitoring of pancreatic cancer. Journal of Nanobiotechnology, 2020. 18(1): p. 150.

14. Kayo, S., et al., A microfluidic device for immuno-aJinity-based separation of mitochondria from cell culture. Lab on a Chip, 2013. 13(22): p. 4467–4475.

15. Takahashi, T., et al., A microfluidic device for isolating intact chromosomes from single mammalian cells and probing their folding stability by controlling solution conditions. Scientific Reports, 2018. 8(1): p. 13684.

16. Lu, H., et al., A Microfabricated Device for Subcellular Organelle Sorting. Analytical Chemistry, 2004. 76(19): p. 5705–5712.

17. Toyama, K., M. Yamada, and M. Seki. Development of microfluidic cell nucleus separator employing rapid chemical treatment. in 2010 International Symposium on Micro-NanoMechatronics and Human Science. 2010.

18. Yamada, M., et al., Millisecond treatment of cells using microfluidic devices via two-step carrier-medium exchange. Lab on a Chip, 2008. 8(5): p. 772–778.

19. Tesauro, C., et al., Isolation of functional mitochondria by inertial microfluidics – a new method to sort intracellular organelles from a small scale biological sample. RSC Advances, 2017. 7(38): p. 23735–23741.

20. Lamanna, J., et al., Digital microfluidic isolation of single cells for -Omics. Nature Communications, 2020. 11(1): p. 5632.

21. Hung, P.-Y., et al., Genomic DNA extraction from whole blood using a digital microfluidic (DMF) platform with magnetic beads. Microsystem Technologies, 2017. 23(2): p. 313–320.

22. Hale, C. and J. Darabi, Magnetophoretic-based microfluidic device for DNA isolation. Biomicrofluidics, 2014. 8(4): p. 044118.

23. Reedy, C.R., et al., Solid phase extraction of DNA from biological samples in a post-based, high surface area poly(methyl methacrylate) (PMMA) microdevice. Lab on a Chip, 2011. 11(9): p. 1603–1611.

24. Lee, C., et al., A Three-Dimensional Printed Inertial Microfluidic Platform for Isolation of Minute Quantities of Vital Mitochondria. Analytical Chemistry, 2022. 94(19): p. 6930–6938.

25. Venkatesan, M., et al., Spatial subcellular organelle networks in single cells. Scientific Reports, 2023. 13(1): p. 5374.

26. Benítez, J.J., et al., Microfluidic extraction, stretching and analysis of human chromosomal DNA from single cells. Lab on a Chip, 2012. 12(22): p. 4848–4854.

27. Wang, X., et al., Microfluidic extraction and stretching of chromosomal DNA from single cell nuclei for DNA fluorescence in situ hybridization. Biomedical Microdevices, 2012. 14(3): p. 443–451.

28. Wood, D.K., et al., Single cell trapping and DNA damage analysis using microwell arrays. Proceedings of the National Academy of Sciences, 2010. 107(22): p. 10008–10013.

29. Hughes, A.J., et al., Single-cell western blotting. Nature Methods, 2014. 11(7): p. 749–755.

30. Kang, C.-C., et al., Single cell–resolution western blotting. Nature Protocols, 2016. 11(8): p. 1508–1530.

31. Yamauchi, K.A. and A.E. Herr, Subcellular western blotting of single cells. Microsystems & Nanoengineering, 2017. 3(1): p. 16079.

32. McCarthy, F.M., et al., DiJerential Detergent Fractionation for Non-electrophoretic Eukaryote Cell Proteomics. Journal of Proteome Research, 2005. 4(2): p. 316–324.

33. Rosàs-Canyelles, E., et al., Assessing heterogeneity among single embryos and single blastomeres using open microfluidic design. Science Advances. 6(17): p. eaay1751.

34. Rosàs-Canyelles, E., et al., Multimodal detection of protein isoforms and nucleic acids from low starting cell numbers. Lab on a Chip, 2021. 21(12): p. 2427–2436.

35. Rosàs-Canyelles, E., et al., Multimodal detection of protein isoforms and nucleic acids from mouse pre-implantation embryos. Nature Protocols, 2021. 16(2): p. 1062–1088.

36. Sandström, N., et al., Live single cell imaging assays in glass microwells produced by laser-induced deep etching. Lab on a Chip, 2022. 22(11): p. 2107–2121.

37. Bromberg, L., et al., Kinetics of Swelling of Polyether-Modified Poly(acrylic acid) Microgels with Permanent and Degradable Cross-Links. Langmuir, 2005. 21(4): p. 1590–1598.

38. Plunkett, K.N., et al., Swelling Kinetics of Disulfide Cross-Linked Microgels. Macromolecules, 2003. 36(11): p. 3960–3966.

39. Wang, Y., et al., Dissolvable Polyacrylamide Beads for High-Throughput Droplet DNA Barcoding. Advanced Science, 2020. 7(8): p. 1903463.

40. Pan, Q., K.A. Yamauchi, and A.E. Herr, Controlling Dispersion during Single-Cell Polyacrylamide-Gel Electrophoresis in Open Microfluidic Devices. Anal Chem, 2018. 90(22): p. 13419–13426.

41. Sheth, S., et al., UV Dose Governs UV-Polymerized Polyacrylamide Hydrogel Modulus. International Journal of Polymer Science, 2017. 2017: p. 5147482.

42. Takemori, A., et al., BAC-DROP: Rapid Digestion of Proteome Fractionated via Dissolvable Polyacrylamide Gel Electrophoresis and Its Application to Bottom-Up Proteomics Workflow. Journal of Proteome Research, 2021. 20(3): p. 1535–1543.

43. Okuno, T. and N. Kondelis, Evaluation of dithiothreitol (DTT) for inactivation of IgM antibodies. J Clin Pathol, 1978. 31(12): p. 1152–5.

44. Gammelgaard, S.K., et al., Direct Ultraviolet Laser-Induced Reduction of Disulfide Bonds in Insulin and Vasopressin. ACS Omega, 2020. 5(14): p. 7962–7968.

45. Wongkongkathep, P., et al., Enhancing Protein Disulfide Bond Cleavage by UV Excitation and Electron Capture Dissociation for Top-Down Mass Spectrometry. Int J Mass Spectrom, 2015. 390: p. 137–145.

46. Corces, M.R., et al., An improved ATAC-seq protocol reduces background and enables interrogation of frozen tissues. Nat Methods, 2017. 14(10): p. 959–962.

